# Reducing transitory starch in cassava impairs photosynthesis and carbon export to storage roots

**DOI:** 10.64898/2026.10.21.760634

**Authors:** Qingting Liu, Christian E. Lamm, Alexander Kaier, Joerg Hofmann, Johannes Schmidt, Tzu Jiu Chiu, Shu-Heng Chang, Wilhelm Gruissem, Uwe Sonnewald, Wolfgang Zierer

## Abstract

Optimizing source–sink relations is a central strategy for improving crop yield. In potato, reducing transitory leaf starch synthesis can redirect carbon to storage organs without impairing growth. To test whether this strategy is transferable to cassava, we suppressed the major small subunit of ADP-GLUCOSE PYROPHOSPHORYLASE, *MeAPS1a*. Transgenic plants were nearly devoid of transitory leaf starch and showed impaired photosynthetic carbon fixation. Under greenhouse conditions, shoots acclimated to the reduced carbon supply and exhibited only moderate biomass losses, and the data indicated reduced carbon supply to developing storage roots. This was reflected by lower leaf sucrose levels, reduced storage root biomass and starch accumulation, and downregulation of carbohydrate-responsive genes, including the major AGPase large subunit *MeAPL3*. Under field conditions, the absence of transitory starch was even more detrimental, causing severe growth inhibition and drastic reductions in both shoot and root biomass. We also found that *MeAPS1a*-derived small interfering RNAs moved systemically from leaves to roots, with lower root siRNA abundance in older, larger plants than in younger, smaller plants. Together, our results demonstrate that transitory starch is indispensable for sustaining carbon flux to cassava storage roots, highlighting species-specific constraints on engineering source–sink relations.

## Introduction

Cassava (*Manihot esculenta* Crantz) is a major staple crop in tropical regions. Its starchy tuberous roots are used to produce a variety of food products consumed by hundreds of millions of people (Howeler et al., 2013). Despite its importance, cassava agriculture faces persistent challenges. Smallholder farmers, who make up most crop producers, especially in sub-Saharan Africa, often lack access to modern agricultural tools, essential resources like fertilizers, and advanced planting materials. Consequently, their cassava yields remain low and are susceptible to a range of biotic and abiotic stresses, including cassava mosaic disease, cassava brown streak disease, drought, and poor soil fertility (Adebayo, 2023; Howeler et al., 2013). Developing cassava varieties with the capacity to produce higher and more stable yields is pivotal to improving food security in cassava-producing regions.

One well-established approach to increasing crop yield is the precise modulation of carbon flux from source leaves to sink tissues. Among the various carbon pools found in leaf tissues, transitory starch has emerged as a particularly compelling resource due to its biochemical plasticity and the substantial natural variation it exhibits across a wide array of species, genotypes, and environmental conditions (Fettke and Fernie, 2024). However, the relationship between transitory starch levels and whole-plant performance, including photosynthetic capacity, carbon export, and storage-organ yield, remains elusive and often unpredictable.

In potatoes, for example, the constitutive suppression of ADP-GLUCOSE PYROPHOSPHORYLASE (AGPase), a pivotal enzyme in starch biosynthesis, has been shown to increase tuber number and fresh weight, while concomitantly reducing tuber starch content (Müller-Röber et al., 1992). More nuanced manipulations demonstrate the system’s flexibility. Reducing AGPase activity by over 90% exclusively in leaves lowered the leaf starch levels to approximately 30% of wild-type levels without altering photosynthetic rates or tuber yield (Leidreiter et al., 1995). Additionally, combining reduced leaf AGPase activity with increased hexose-phosphate and adenosine triphosphate import into tubers (through *GLUCOSE-6-PHOSPHATE/PHOSPHATE TRANSLOCATOR* and *ADENYLATE/NUCLEOTIDE TRANSPORTER* overexpression) increased tuber starch content and greenhouse biomass (Jonik et al., 2012; Kammerer et al., 1998; Kampfenkel, 1995; Zhang et al., 2008). Together, these studies suggest that elevated levels of transient starch are not necessarily required to sustain photosynthesis or tuber yield in potato. In contrast, *Arabidopsis thaliana* exhibits a strong dependence on transitory starch. The *pgm1* mutant lacks PHOSPHOGLUCOSEMUTASE (PGM), a key enzyme in starch synthesis. This mutant exhibit severely impaired growth, almost no leaf starch, and elevated levels of soluble sugars (Caspar et al., 1985; Kofler et al., 2000).

Interestingly, knocking down PGM isoforms in hybrid aspen resulted in the complete elimination of leaf starch and reduced photosynthetic capacity. However, this did not affect tree growth or wood density under greenhouse conditions (Wang et al., 2022). Poplar is believed to be a predominantly symplasmic phloem loader, with a high number of plasmodesmata connections that facilitate diffusive sugar allocation (Rennie and Turgeon, 2009; Zhang et al., 2014). Cassava, a perennial woody shrub, exhibits physiological and anatomical characteristics typical of symplasmic loaders. These characteristics include elevated levels of leaf sucrose and clusters of plasmodesmata at the interface between companion and bundle sheath cells (Rüscher et al., 2024).

Together, these studies reveal the remarkable flexibility and species-specific diversity in how plants partition carbon between source and sink tissues. It is hypothesized that reducing transitory starch synthesis in certain species may result in a greater proportion of photoassimilates being directed toward storage tissues without compromising growth or carbon assimilation. The potential of analogous strategies to enhance carbon allocation and yield in cassava remains to be elucidated.

In this study, we examined the role of transitory leaf starch in cassava by suppressing the gene that encodes the major catalytically active AGPase small subunit (*MeAPS1a*). We used a green-tissue-specific promoter to express a *MeAPS1a* RNA interference construct. This approach reduced transitory starch in autotrophic tissues, enabling us to evaluate the impact of reduced leaf starch on photosynthetic capacity, diurnal carbon dynamics, sucrose export, and storage root development under both greenhouse and field conditions. We also evaluated whether the observed reduction in *MeAPS1a* mRNA observed in storage root tissues resulted from the movement of silencing signals from the shoot to the root or was rather a metabolic consequence of the source manipulation.

Our results demonstrate that, unlike poplar or potato, cassava depends more heavily on transitory starch to sustain photosynthesis and maintain carbon flux into storage roots. Loss of leaf starch significantly reduced photosynthetic CO₂ fixation and suppressed sucrose export. It also substantially impaired storage root formation, particularly under field conditions. Although we observed long-distance movement of *MeAPS1a*-derived siRNAs, their low accumulation in mature roots was insufficient to explain the severe sink defects. Taken together, these findings elucidate the critical necessity of transitory starch for the source–sink equilibrium in cassava, thereby emphasizing constraints that must be considered when engineering cassava to enhance yield.

## Material and Methods

### Construct cloning

The *MeAPS1a* RNAi construct was created through Golden Gate cloning. Level 1 of the plasmid was created by combining the *pAtCAB1* promoter fragment, an 18-bp 5′ UTR linker sequence (<u>TACT</u>GAATTCGGATTCTCTAGA<u>AATG</u>), the *MeAPS1a* short hairpin construct, and the *SIATPase* terminator (Engler et al., 2014). The linker sequence and the *MeAPS1a* short hairpin construct were obtained from a DNA synthesis company (Metabion). The short hairpin construct was designed using a 243 bp *MeAPS1a* targeting sequence in the forward direction, followed by a 93 bp linker sequence, and the 243 bp reverse *MeAPS1a* complement sequence. The alignment of the target sequence to the *MeAPS1a* transcript is shown in **Fig. S5**. Level 2 of the plasmid was created by combining the level 1 plasmid with the p134GG cassava transformation plasmid (Mehdi et al., 2019). The plasmid is shown in **Fig. S5**.

### Cassava transformation and tissue culture

The transgenic *MeAPS1a* RNAi plants were produced via *Agrobacterium tumefaciens*-mediated tissue culture transformation of the cassava cultivar TMS60444 as described previously (Bull et al., 2009). The plantlets were maintained in tissue culture jars containing Murashige and Skoog medium with vitamins (M0222, Duchefa Biochemie, Haarlem, The Netherlands), supplemented with sucrose (20 g/l) and CuSO_4_ (2 µM). The plantlets were grown in a growth cabinet under white light (approximately. 120 µmol/m²/s) with a 16/8-hour day/night cycle at a temperature of 28°/26°C (day/night). Plantlets were renewed regularly through stem cuttings (approximately every six weeks).

### Phylogenetic tree construction

The six Arabidopsis APL and APS genes were selected using keyword search on TAIR (Reiser et al., 2024). Corresponding orthologs were obtained from the AM560-2 (v8.1) (Lyons et al., 2022) and TME204 cassava genome assemblies (Qi et al., 2022), and the reference genome of *Populus trichocarpa* (v4) (Tuskan et al., 2006) based on BLASTN similarity to the Arabidopsis coding sequences (e < 1e^−5^) and the presence of the three Pfam domains found in the protein sequences (PF25247.2, PF00483.30, and PF24894.2). Among equally ranking BLAST results, the longest transcript was selected. Full-length protein sequences were aligned using the auto option in MAFFT (v7.490) (Katoh, 2002). Maximum likelihood trees were generated using the phangorn package (v2.12.1) (Schliep, 2011) in R (v4.5.3). The JTT+G (4) model was chosen as the best-fitting model by the implemented modelTest function, with a Bayesian Information Criterion (BIC) of 23,518.4. Maximum-likelihood trees were then constructed using this model and the bootstrap.pml function was used with 100 bootstrap iterations and Nearest Neighbor Interchange (NNI) topology optimization.

### Determination of chlorophyll

Chlorophyll a/b levels were determined following the method (Lichtenthaler, 1987). Chlorophyll a and b were calculated as follows:

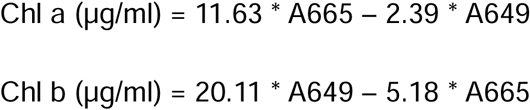

### Greenhouse growth conditions

Cassava plants were grown in a greenhouse in Erlangen, Bavaria, Germany. The conditions were set to 30°/28°C (16-hour day/ 8-hour night), 60% relative humidity, and 180 μmol photons m^−2^ s^−1^. The tissue culture plantlets were carefully potted in a soil mixture of standard soil (type ED73, Einheitserde Patzer; Sinntal-Altengronau, Germany) and 10% fine sand. The soil mixture was initially watered with Gnatrol®SC (Biofa GmbH, Münsingen, Germany) to suppress the growth of fungus gnat larvae. The freshly potted plants were placed in a high-humidity chamber for one week, after which they were progressively acclimated to lower humidity over the following week. After two weeks, the plants were grown in the greenhouse and watered daily.

### Grafting experiments

The cassava plants were grown for four weeks after planting into the soil before grafting. All mature leaves were removed, and the shoot was cut close to the base of the stem, leaving approximately 25% of the stem on the rootstock. The shoot stem was then cut into a V-shape and grafted onto a vertical V-shaped cut rootstock. The scion and rootstock were fused and tightly secured with parafilm. The grafted plants were grown under high humidity for three days and were then adjusted to lower humidity. After one week, the plants were transferred to the greenhouse.

### Confined field trial

Confined field trials (CFTs) were conducted in 2022 at the National Chung Hsing University (NCHU) Experimental Station in Taichung, Taiwan (latitude 24°4’41.50“N; longitude 120°42’56.26“E). The field trial and harvest were conducted as described previously (Zierer et al., 2025).

### Gas exchange measurements

A LI-COR 6800 Portable Photosynthesis System was used for the gas exchange measurements, which were taken from 10:00 to 17:00 for the plants of the indicated age. The following gas exchange parameters were recorded under the given environmental conditions: carbon assimilation (A), intercellular CO2 concentration (Ci), and stomatal conductance (gsw). Flow: 600 μmol/s; H_2_O: 1.3-1.5 kPa VPD (same for all plants in one experiment); CO_2_ sequence: 400, 300, 200, 100, 50, 0, 400, 400, 600, 800, 1000, 1200, 1400, 1600 μmol/mol; temperature: Tair 28-30 °C (same for all plants in one experiment); fan speed: 10,000 rpm; light: 1,000-15,000 μmol m^−2^ /s^−2^, adapted from Ruiz-Vera (2021). The stability criteria for logging are: gsw: slope limit 0.1; stdev limit −0.1; A: slope limit 1.5, stdev limit 0.5.

Gas exchange measurements were analyzed with a Python (v 3.8) script that fitted the experimental data to photosynthesis models described by (S von Caemmerer, 2000) via non-linear least-squares minimization implemented in the LMFIT package (Newville et al., 2014). As described in (Ruiz-Vera et al., 2021), each measurement was divided into two groups to model the RuBP-limited and export-limited states separately.

### RNA extraction and gene expression analysis

RNA extractions were performed according to the instructions included with the Spectrum Plant Total RNA kit (Sigma, STRN250). DNase digestion and cDNA synthesis were performed using the RevertAid First Strand cDNA Synthesis Kit (Thermofisher, K1621). The GoTaq Master Mix (Promega, A600A) was used for qPCR with the primers listed in **Suppl. Table 1.** The AriaMx Real-Time PCR System (Agilent Technologies) was used for the qPCR reactions with the following thermal profile: 95 °C for 2 min, [95 °C for 5 sec, 60 °C for 30 sec] x 40 cycles, 95 °C for 30 sec, 55 °C for 30 sec, 95 °C for 30 sec. The expression levels of the examined genes were normalized to *MeGAPDH* or *MePP2A*.

### Starch staining

Leaf discs were destained in 95% ethanol at 80°C until colorless and then rinsed with water and stained with an iodine solution (0.23% [w/v] KI, 0.13% [w/v] I_2_).

### Determination of sugars, starch, and phosphorylated intermediates

The extraction of soluble carbohydrates was performed using the perchloric acid method previously described by (Rosado-Souza et al., 2019). The same report describes the extraction and hydrolysis of starch. The quantification of extracted sugars and hydrolyzed starch was performed using an NAD^+^-coupled enzymatic assay as previously described by (Stitt et al., 1989). For this analysis, the absorbance of NADPH was measured at 340 nm using a photometer (Tecan Infinite M Nano, Männedorf, Switzerland). The sugar concentration was then calculated according to the Lambert-Beer law. The extraction and measurement of phosphorylated intermediates were performed as previously described (Rosado-Souza *et al*., 2019).

### Determination of AGPase activity

The frozen plant material was ground with liquid nitrogen. Crude proteins were extracted from approximately 100 mg of the fine powder using an extraction buffer [50 mM HEPES-KOH, pH 7.5, and 5mM MgCl_2_] followed by centrifuging at 4 °C for 10 min. The resulting supernatant was used for both the Bradford assay, which determines the protein concentration, and the enzyme assay.

The Bradford assay was performed according to the instructions provided with the reagents (Bio-Rad, #5000006).

The AGPase assay was adapted from Tiessen *et al*. (2002). The reaction buffer contains: 50 mM HEPES-KOH, pH 7.5; 5 mM MgCl_2_; 10 μM glucose-1,6-bisphosphate; 0.6 mM NAD^+^; 2.5 mM Na-PPi; 1 U/mL phosphoglucomutase; 2.5 U/mL glucose-6-phosphate dehydrogenase; 1 mM ADP-glucose and 5 mM DTT. The reaction was initiated by adding 20 μL of crude protein and 200 μL of reaction buffer to a 96-well plate. The absorbance change at 340 nm was monitored with a photometer. Rates were calculated using the Beer-Lambert law with ε_340_ = 6.22 mM^−1^ cm^−1^ and an effective path length of 0.6 cm (reaction volume 220 µL). Activities were reported as extractable activities, and values were normalized to fresh weight and soluble protein.

### Small RNA sequencing and analysis

Total RNA was extracted from the designated organs/tissues using a CTAB-based protocol. Plant materials were ground in liquid nitrogen and then added to 1 mL of CTAB buffer solution containing 2% CTAB; 2% PVP-40, 100 mM Tris-HCl, pH 8.0, 25 mM EDTA, 2 M NaCl, 0.5 g/L spermidine; 2% beta-mercaptoethanol, and adjusted to the given volume with DEPC-pre-treated water. The sample was thoroughly mixed and then incubated at 50 °C for 15 min with occasional shaking, followed by centrifugation for 5 min at 25000 g, and 900 μL of the supernatant was transferred to a clean 2 mL tube. 900 μL of a chloroform: isoamyl alcohol mixture (24:1) was added, mixed with the supernatant, and centrifuged at 25000g for 10 min. 800 μL of the supernatant was transferred to a fresh tube, an equal volume of chloroform: isoamyl alcohol (24:1) was added, and the centrifugation was repeated. 600 μL of supernatant was transferred to a fresh 1.5 mL tube. 900 μL of pre-chilled 100% ethanol was added and the mixture incubated at −20 °C for one h or overnight. The mixture was centrifuged for 30 min at full speed and 4 °C. The supernatant was then removed, and the pellet was washed with 80% ethanol, followed by centrifugation for 5 min. The pellet was vacuum-dried and dissolved in 100 μL DEPC-treated water.

Novogene handled the quality control of the extracted RNA, library preparation, sequencing, and data quality control. Alignment was conducted against the full-length mRNA of *MeAPS1a*. This was done using Bowtie (v1.2.3) (Langmead et al., 2009) with the flags -no-unal, -S, and -v 0 to retrieve a SAM file containing only sRNA reads mapping to the full-length mRNA of *MeAPS1a* without any mismatches. Subsequent quantification of siRNA levels was performed using a custom R script and the Biostrings (v2.76.0) (H. Pagès, 2017) and tidyverse (v2.0.0) (Wickham et al., 2019) libraries.

## Results

### Silencing of *MeAPS1a* impairs leaf starch synthesis and photosynthetic capacity

We used BLAST analysis against the AM560 v8.1 (Bredeson et al., 2016) and TME204 cassava genomes (Qi et al., 2022) to identify the cassava homologs of the small (*APS*) and large (*APL*) subunit genes encoding cassava AGPase using the annotated *Arabidopsis thaliana* and *Populus trichocarpa* genes. Nine cassava genes were identified, and these genes, along with those from the other two species, were used to construct a phylogenetic tree **(Fig. S1A)**. The tree is divided into five clades: two *APS* clades and three *APL* clades. The cassava *APS* genes were designated as *MeAPS1a* (*Manes.12G067900*), *MeAPS1b* (*Manes.13G058900*), *MeAPS2* (*Manes.05G208000*), while the APL genes were designated as *MeAPL1a* (*Manes.03G182100*), *MeAPL1b*(*Manes.15G025400*), *MeAPL2a* (*Manes.01G236700*), *MeAPL2b* (*Manes.18G019650*), and *MeAPL3* (*Manes.11G085500*). Next, we analyzed mRNA levels in ten distinct cassava tissues for the *MeAPS* and *MeAPL* genes **(Fig. S1B-I)** using a comprehensive cassava RNA-Seq dataset (Rüscher *et al*., 2024). This analysis revealed that *MeAPS1a* is the primary gene variant among the three *MeAPS* genes, with expression levels two orders of magnitude higher than those of *MeAPS1b* and *MeAPS2*. *MeAPS1a* expression was highest in source leaves and storage roots **(Fig. S1B)**. Among the *MeAPL* genes, *MeAPL1a* **(Fig. S1E)**, *MeAPL1b* **(Fig. S1F)**, and *MeAPL3* **(Fig. S1I)** exhibited the highest expression levels. *MeAPL1a* and *MeAPL1b* were rather specific to source leaves, while *MeAPL3* was rather specific to storage roots. The role of *MeAPL3* in storage root starch synthesis was previously demonstrated using transient and stable gene silencing methods (Beyene et al., 2022). Furthermore, *MeAPL3* expression significantly increased during storage root bulking **(Fig. S1I)**. Notably, *MeAPS1a* expression substantially increased during storage root bulking as well, analogous to that of *MeAPL3* (“DB” in **Figs. S1B** and **Fig. S1I**). This suggests that *MeAPS1a* is the predominant small subunit in storage root starch synthesis. In the source leaves, *MeAPS1a*, *MeAPL1a*, and *MeAPL1b* likely represent the key *MeAPS* and *MeAPL* gene variants for transitory starch synthesis.

To understand carbon reallocation from source leaves to storage roots, we targeted the catalytically active small subunit of AGPase for reducing transient leaf starch levels, similar to previous strategies in potato (Jonik et al., 2012; Leidreiter et al., 1995). We designed a short hairpin RNA (shRNA) construct against *MeAPS1a*, which is the *MeAPS* gene variant most highly expressed in cassava leaves **(Fig. 1A)**. The *pAtCAB1* (*At1g29930*) promoter, which is active only in green cassava tissues (Zierer et al., 2022) was used to specifically target the expression of the *MeAPS1a* shRNA to autotrophic tissues to silence *MeAPS1a*. The construct was transformed into the cassava genotype TMS60444, and 11 independent transgene-positive lines were initially recovered. A notable reduction in transitory leaf starch was observed in all 11 lines as evidenced by iodine staining **(Fig. S1J)**. We selected the transgenic lines RNAi_61, 154, 165, and 171 for a comprehensive analysis.

**Fig. 1.**
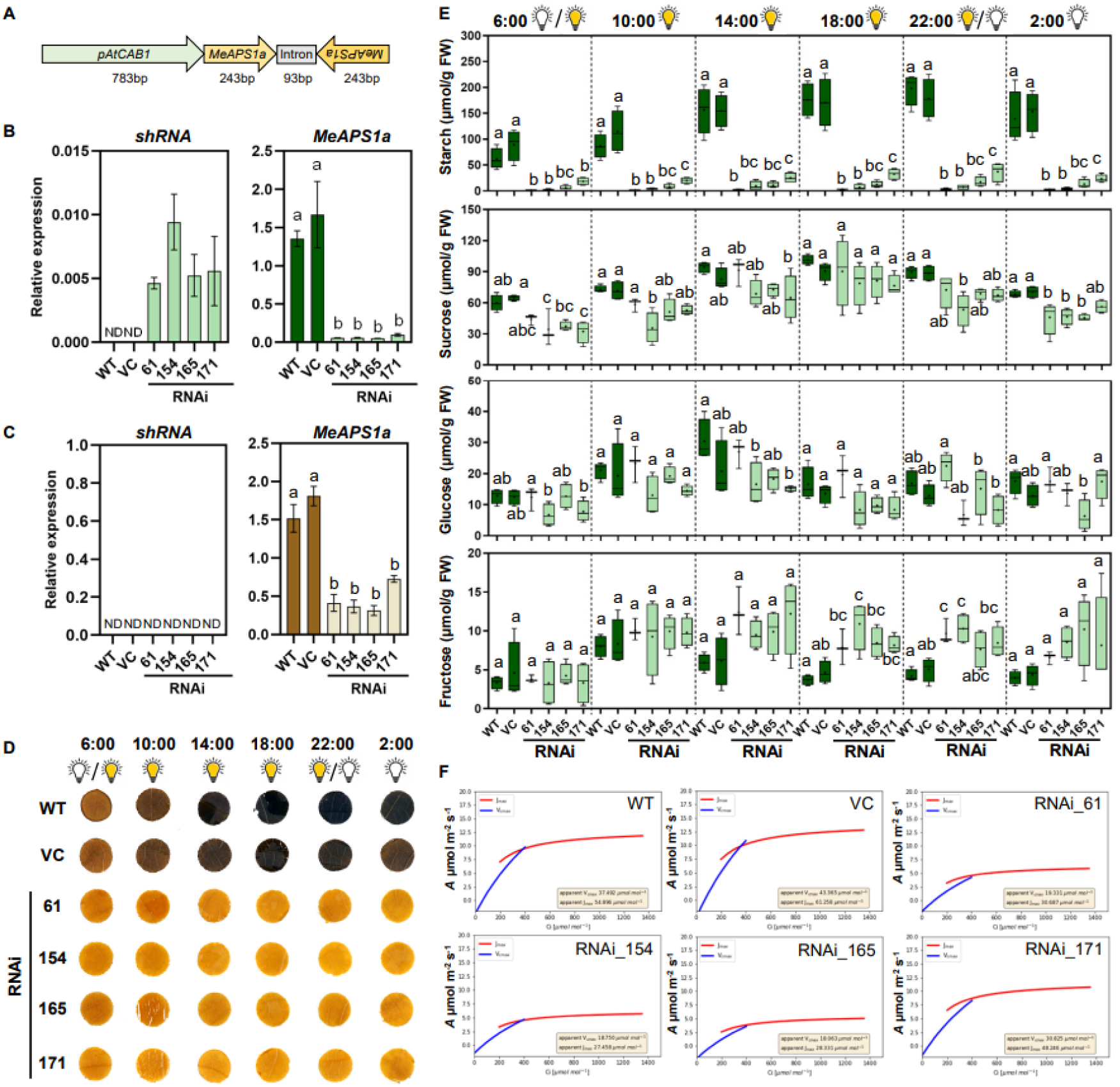
RNA interference represses *MeAPS1a* expression and impairs leaf starch synthesis. **A**) Illustration of the shRNA transformation cassette utilized in the downregulation of *MeAPS1a* expression. **B**) The shRNA and *MeAPS1a* transcript levels in source leaves. ND = not detected. N = 4. **C**) The shRNA and the *MeAPS1a* transcript abundance in storage roots. ND = not detected. **D**) Iodine staining of cassava source leaf discs from wildtype (WT), vector control (VC), *MeAPS1a*_RNAi_61, 154, 165, and 171 lines throughout the light (6:00-22:00) and dark period (22:00-6:00). **E**) Concentrations of starch, sucrose, glucose, and fructose measured in WT, VC, and *MeAPS1a*_RNAi plant lines during the light-dark cycle. The letters in each graph indicate significant differences based on one-way ANOVA analysis with Tukey’s post hoc test (*p*<0.05). **F**) Assimilation/intercellular CO_2_ (A/Ci) response curves of WT, VC, and *MeAPS1a*_RNAi plant lines. All plants tested were 8 weeks old.

To validate the specific silencing of *MeAPS1a*, the expression levels of shRNA and *MeAPS1a* were subsequently quantified in source leaves and storage roots **(Fig. 1B, C)**. As expected, the shRNA was detectable in the source leaves of the *MeAPS1a* RNAi lines and *MeAPS1a* mRNA levels were significantly reduced **(Fig. 1B)**. Interestingly, *MeAPS1a* mRNA levels were also considerably reduced in the storage roots, despite the use of green tissue-specific promoter and the absence of detectable local shRNA expression **(Fig. 1C)**, suggesting a mobile effect from the source (leaf) to the sink (storage root).

To confirm the specificity of the *MeAPS1a* RNAi, we measured the mRNA levels of all other *MeAPS* and *MeAPL* genes in the representative vector control (VC) and the four selected RNAi lines **(Fig. S1K)**. In source leaves, the expression levels of the other tested genes remained unchanged, except for *MeAPL3.* Previous studies have reported that *MeAPL3* has a primary role in storage root starch synthesis (Beyene *et al.,* 2022). *MeAPL3* exhibited significantly decreased expression levels in the four RNAi lines (**Fig. S1H**)*. MeAPS1a* shares over 90% DNA sequence identity with *MeAPS1b,* whose expression remained unchanged in source leaves. However, *MeAPS1a* shares about 60% identity with *MeAPL3*. Therefore, *MeAPL3* is unlikely to be directly repressed by RNAi derived from the *MeAPS1a* shRNA. However, it is plausible that *MeAPL3* expression is co-regulated with *MeAPS1a* or affected metabolically.

To assess the effect of the significant decrease in *MeAPS1a* mRNA levels, we first determined leaf starch levels throughout the day and night using iodine staining **(Fig. 1D)** and an enzymatic assay **(Fig. 1E)**. The starch content of the RNAi lines decreased by 80% (line 171) to 98% (line 61) compared to the control plants by the end of the light phase (22:00). Unlike potato, sucrose, which is used for carbon export to sink organs, did not increase in response to decreased leaf starch levels in cassava (Leidreiter *et al*., 1995; Jonik *et al*., 2012). In fact, sucrose levels were significantly lower in the *MeAPS1a* RNAi lines, particularly at night. This decline is likely due to an inability to convert sufficient levels of transitory starch to sucrose and increased sucrose catabolism **(Fig.1E)**. After the onset of light (6:00), leaf sucrose levels were generally lower in the *MeAPS1a* RNAi lines than in the control plants.

However, sucrose levels gradually increased, reaching control levels only after 12 hours of light exposure **(Fig. 1E)**. While the steady-state glucose levels of *MeAPS1a* RNAi plants were similar to those of control plants, fructose levels were increased overall **(Fig. 1E).** Trehalose-6-phosphate (T6P) and sucrose-6-phosphate (S6P) levels were decreased in the leaves of *MeAPS1a* RNAi plants during the day (**Fig. S2A**). Since T6P functions as a signaling molecule for the sucrose status and S6P is the biosynthetic precursor of sucrose (Figueroa and Lunn, 2016; Lunn et al., 2006), this observation is consistent with reduced sucrose levels. Furthermore, the decline in multiple phosphorylated metabolites, including TCA cycle intermediates, suggests that leaf metabolism in *MeAPS1a* RNAi plants was severely impacted by a carbon shortage **(Fig. S2B-D)**.

To evaluate the photosynthetic capacity of the *MeAPS1a* RNAi plants, we conducted CO_2_-response assays using at least three individual plants for each transgenic event and calculated fitting curves **(Fig. 1F).** Comparing the RNAi lines to the wild type (WT) and VC plants revealed a significant reduction in photosynthetic capacity, with a decline of over 50% in three out of four RNAi lines. Interestingly, *MeAPS1a* mRNA levels were less reduced in line RNAi_171 compared to the other three lines. The RNAi_171 plants exhibited the greatest retention of leaf starch and the least decrease in photosynthetic capacity among the *MeAPS1a* RNAi lines. These results suggest that minimal leaf starch synthesis is sufficient to circumvent severe limitations in triose phosphate utilization.

In summary, the silencing of *MeAPS1a* in leaves resulted in nearly complete loss of transitory starch and significantly impaired photosynthetic carbon fixation. Carbohydrate data also suggest a reduction in sucrose export to sink organs in response to *MeAPS1a* silencing, due to lower sucrose levels throughout the day and night.

### Reducing leaf *MeAPS1a* expression reduces storage root biomass

The tissue-cultured *MeAPS1a* RNAi plantlets experienced slower growth in the climate chamber and after being transplanted to soil in the greenhouse. A comparison of the stem height and lower stem diameter of the more mature *MeAPS1a* RNAi plants revealed no significant differences compared to the control plants **(Fig. 2A-C)**. However, while the control plants had an average of ten leaves, *MeAPS1a* RNAi plants had an average of only eight leaves **(Fig. S2E)**. Photometric measurements of *MeAPS1a* RNAi plant leaves revealed reduced levels of chlorophyll a and b **(Fig. 2D-E)**. Interestingly, the *MeAPS1a* RNAi plants had significantly larger leaves than the control plants **(Fig. 2F; S2A)**, suggesting a compensatory effect at the organ level, possibly in response to the reduced photosynthetic performance per area **(Fig. 1F)**. The levels of the phytohormones auxin (IAA) and the cytokinin N6A, zeatin, and zeatin riboside were significantly higher in the *MeAPS1a* RNAi plants than in the control plants **(Fig. S2F-G)**. Conversely, jasmonic acid and salicylic acid levels were reduced in the RNAi plants **(Fig. S2H-I)**. Abscisic acid and gibberellic acid concentrations did not differ between RNAi and control plants **(Fig. S2G-K)**. However, both fresh weight **(Fig. 2G)** and dry shoot weight **(Fig. 2H)** were significantly reduced in *MeAPS1a* RNAi plants compared to control plants. Meanwhile, the relative dry matter content remained at the level observed in control plants **(Fig. 2I)**.

**Fig. 2.**
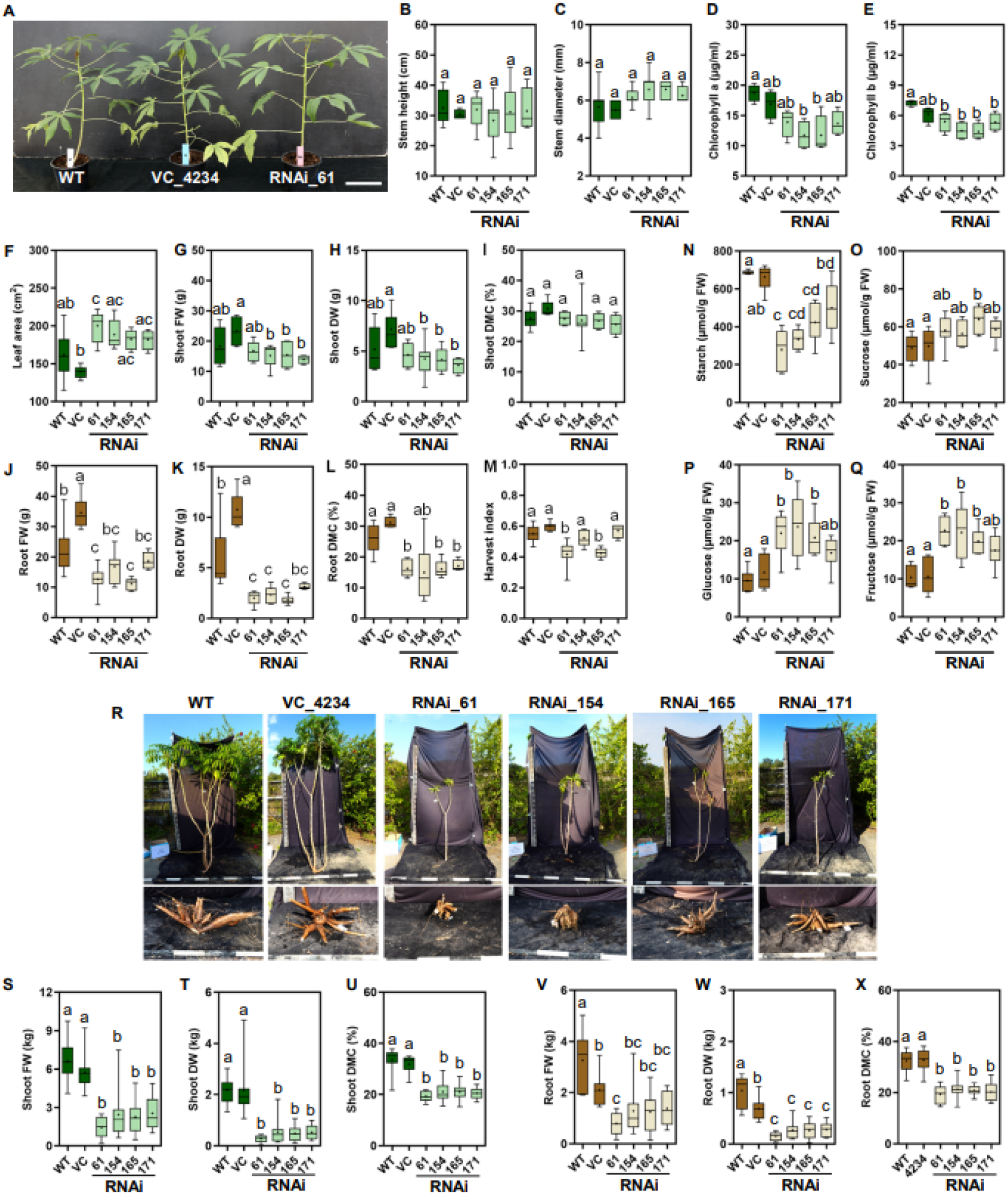
Phenotypes of *MeAPS1a*_RNAi plants under greenhouse and field conditions. **A**) Representative phenotypes of WT, VC, and *MeAPS1a*_RNAi plant lines are presented in greenhouse conditions. Scale bar = 10cm. **B-C**) Plant stem height and lower stem diameter of WT, VC, and *MeAPS1a*_RNAi plant lines. **D-E**) The concentration of chlorophyll a and b in leaves. **F**) Leaf area (the 1^st^ to 5^th^ source leaves from the shoot apex). **G-I**) Shoot fresh weight (FW), shoot dry weight (DW), and dry matter content (DMC) of WT, VC, and *MeAPS1a* RNAi plant lines. **J-L**) Storage root FW, DW, and DMC. **M**) Harvest index. **B-M**) N = 4 to 10 plants for each line. **N-Q**) Starch, sucrose, glucose, and fructose content of the storage roots. N = 5 to 8 plants for each line. **B-Q**) The letters in each graph indicate the significance between plant lines based on one-way ANOVA analysis with Tukey’s post hoc test (*p*<0.05). **R**) Plant photos were taken at harvest after 8 months of field growth. **S-X)** Shoot FW, shoot DW, shoot DMC, root FW, root DW, root DMC of WT, VC, and RNAi plants. N = 8-10. Letters indicate the significant difference between the control plants (WT and VC) and RNAi plants based on one-way ANOVA analysis with Tukey’s post hoc test (*p*<0.05).

Although aboveground growth was less affected in the *MeAPS1* RNAi lines than in the control plants in the greenhouse, storage root biomass was significantly reduced. The results showed a substantial decline in storage root fresh weight **(Fig. 2J)** and dry weight **(Fig. 2K)**. Consequently, both the relative dry matter content of the storage root **(Fig. 2L)** and the plant harvest index **(Fig. 2M)** were decreased. These reduced agronomic parameters of the storage root are consistent with reduced carbon export from the *MeAPS1a* RNAi source leaves.

Storage root starch levels were significantly lower in *MeAPS1a* RNAi plants than in controls. The reduction pattern followed leaf starch levels and photosynthetic capacity (*i.e.*, line 171 displayed the least reduction; **Fig. 2N**). The small increase in storage root sucrose levels was significant in only one out of four lines **(Fig. 2O)**. Storage root glucose **(Fig. 2P)** and fructose **(Fig. 2Q)** levels increased, likely due to increased sucrose cleavage by invertases. This process has been associated with low starch levels in different cassava genotypes (Lamm et al., 2023). Collectively, the data suggest that compromised transitory starch synthesis in cassava severely impairs photosynthetic capacity, although the plants maintain the capacity for reasonable shoot growth in a greenhouse environment. However, the substantial decrease in sucrose availability during the day and the sucrose derived from starch degradation at night suggest reduced transport to storage roots. This reduction in transport is likely the cause of the significant reductions in storage root biomass production observed.

Based on the results from the greenhouse-grown *MeAPS1a* RNAi plants, we hypothesized that the growth penalty from transitory leaf starch reduction would be more pronounced under field conditions. To test this hypothesis, we conducted a field trial with the *MeAPS1a* RNAi plants at the National Chung-Hsing University (NCHU) Experimental Station, Taichung, Taiwan. At the time of harvest, the growth of the *MeAPS1a* RNAi plants was significantly reduced compared to the control plants **(Fig. 2R)**. The relative reductions in shoot fresh weight **(Fig. 2S),** shoot dry weight **(Fig. 2T),** and shoot dry matter content **(Fig. 2U)** were drastically greater than in the greenhouse. However, the reductions in storage root fresh weight (**Fig. 2V**), storage root dry weight (**Fig. 2W**), and storage root dry matter content (**Fig. 2X**) were similar to those observed in the greenhouse. Together, these data demonstrate that, although cassava can maintain its growth capacity with reduced levels of transitory starch, its agronomic performance and storage root production are greatly diminished.

### *MeAPS1a* silencing in storage roots was detected only in small grafts

Due to the significant decrease in *MeAPS1a* expression observed in the storage roots, we hypothesized that a mobile effect was occurring from source to sink, potentially via small RNAs. To determine if small RNAs processed from the *MeAPS1*a shRNA moved from the shoot to the root and silenced *MeAPS1a*, we grafted a vector control line (VC) and a *MeAPS1a* RNAi line (61) reversibly as scion x rootstock (VCxVC, VCx61, 61×61, and 61xVC). Two groups were designated as small pot grafts (SG) and large pot grafts (LG). All grafts were initially cultivated in small pots. At 29 days after grafting (DAG), the LG plants were transferred to larger pots to allow continued growth. The SG plants remained in small pots. The SG and LG plants were sampled at 44 and 127 DAG, respectively, allowing comparison of silencing at different plant sizes and developmental stages (**Fig. 3A**). The SG or LG plants exhibited similar growth patterns regardless of the graft combination **(Fig. 3B).** We determined the transcriptional activity of *MeAPS1a* in the storage roots and found that *MeAPS1a* expression was downregulated at 44 DAG in SG plants 61×61 and VCx61, but not in SG plants VCxVC and 61xVC **(Fig. 3C)**. The absence of *MeAPS1a* downregulation in 61xVC was unexpected. A second reciprocal grafting experiment, conducted only in small pots using *MeAPS1a* RNAi line 154 and WT plants, revealed *MeAPS1a* downregulation in the storage roots of both 154xWT and WTx154 grafts **(Fig. S3A-B)**. Thus, repression was also observed when the RNAi tissue was confined to the rootstock. None of the LG plants showed repressed *MeAPS1a* expression in the storage root at 127 DAG, regardless of core or peel **(Fig. 3C)**.

**Fig. 3.**
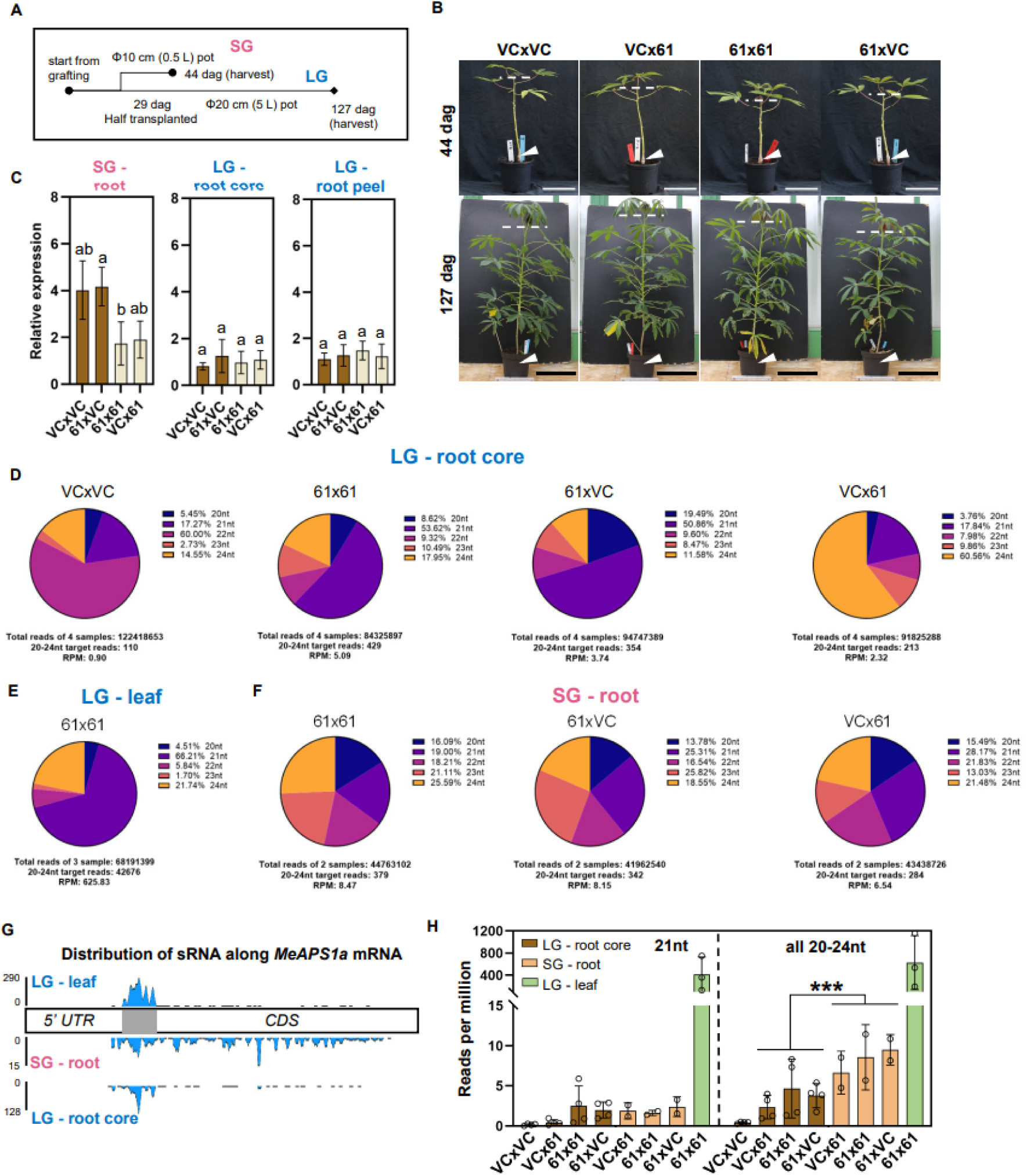
Systemic movement of *MeAPS1a*-derived siRNAs and stage-dependent root *MeAPS1a* repression. **A)** Schematic explanation of grafted plants designated as small pot grafts (SG; sampled at 44 days after grafting) and large pot grafts (LG; sampled at 127 days after grafting). dag: days after grafting. **B)** Photographs of grafted plants at the indicated age in greenhouse conditions. The white triangles indicate the junction of scion and rootstock. The white dashed lines mark the stem apex point. Scale bars = 10cm (white) / 25cm (black). **C**) Transcriptional expression of *MeAPS1a* in the storage root of SG and LG plants. The letters in each graph indicate significant differences based on one-way ANOVA analysis with Tukey’s post hoc test (*p*<0.05). **D-F**) Abundance and proportion of 20-24nt target siRNAs in the designated grafting combination and tissues. RPM, reads per million. **G**) The distribution of siRNA along the *MeAPS1a* mRNA in three tissues. The y-axis numbers indicate the coverage for each nucleotide. The gray box represents the sequence used for shRNA construction. The unique reads of the designated tissue were collected from all tested samples, regardless of grafting combination. **H**) Abundance of siRNAs in each grafting combination. Asterisks indicate the significant difference between the LG storage root core and SG root samples. A Student’s *t*-test was conducted. ***, *p*<0.005.

To assess the presence of *MeAPS1a*-derived small RNAs, we extracted total RNA from leaves and storage roots for small RNA sequencing (sRNA-Seq). This allowed us to determine whether the observed differences in *MeAPS1a* expression could be attributed to differences in siRNA abundance. Since starch synthesis in the cassava storage roots occurs predominantly in the core tissue, we first evaluated *MeAPS1a* target reads in four storage root cores replicates of each LG grafting combination **(Fig. 3D)**. The LG VCxVC combination, which served as a negative control, yielded 110 20–24 nt target reads out of 122 million total reads. These reads perfectly matched the *MeAPS1a* transcript and corresponded to 0.9 reads per million (RPM). Given their very low abundance, these reads likely reflect background RNA turnover products or endogenous RNA surveillance pathways, including occasional RDR-dependent small RNA generation during mRNA decay, rather than specific silencing activity (Tsuzuki et al, 2017). LG 61×61 plants exhibited the highest number of *MeAPS1a*-related siRNAs in the storage root core compared to the other LG plants (429 reads, 5.09 RPM), while LG 61xVC plants had 354 reads (3.74 RPM). The 21 nt siRNA class is predominant in these samples. In contrast, LG VCx61 plants had only 213 target 20-24 nt siRNAs reads (2.32 RPM) and the 21nt siRNA class was not predominant.

Over 68 million total reads were obtained from three leaf samples of LG 61×61 plants to determine the origin of the siRNAs **(Fig. 3E)**. A total of 42,676 target reads of 20-24 nt were obtained, accounting for 625.83 RPM. The 21nt siRNA is the predominant class, accounting for approximately 65% of the total target siRNAs.

The unpeeled root of SG plants was also used for sRNA-Seq. Target 20-24 nt siRNAs exhibited comparable levels in 61×61 and 61xVC plants. However, the RPM was more than doubled in SG 61xVC roots compared to the LG 61xVC storage root core **(Fig. 3F)**. SG VCx61 plants had fewer root siRNAs than 61×61 and 61xVC roots, though more than were found in the storage root core of LG VCx61 plants. Notably, SG roots have an almost equal distribution of the five siRNA classes. The 21-nt siRNA class is no longer predominant in SG 61×61 roots, suggesting a distinct siRNA production mechanism in small cassava roots.

To understand their distribution along the sequence, we aligned the unique 20-24 nt siRNA sequences with the *MeAPS1a* mRNA **(Fig. 3G)**. Of the 42,676 target reads from the three LG leaf samples, 1,516 unique 20-24 nt siRNAs were identified. Of these, 1,469 (96.9%) mapped to the sequence used for shRNA construction and are referred to as primary siRNAs. Additionally, 47 (3.1%) unique siRNAs mapped evenly across the *MeAPS1a* transcript **(Fig. 3G)**. A similar distribution pattern was also observed in the LG storage root core, which contained only 366 unique siRNAs. Interestingly, the 577 unique siRNA sequences from all the SG root samples tested were evenly distributed across the *MeAPS1a* mRNA sequence, as opposed to the unique siRNA sequences from the LG leaves and storage root cores. In this case, only 125 (34%) of the unique siRNA sequences mapped to the shRNA sequence, while the remaining sequences are consistent with secondary siRNAs production, potentially involving an RDR6-dependent mechanism (Klahre et al., 2002; Taochy et al., 2017). The molecular basis for this difference is currently unknown.

Collectively, SG plants accumulated significantly higher levels of the target 20-24 nt siRNAs in the recipient roots than LG plants **(Fig. 3H)**. However, this difference between grafts was modest compared with the much larger difference observed between source leaves and roots. The distribution of siRNA outside the hairpin-targeted region in SG roots is consistent with secondary siRNA production, which could contribute to the repression observed in these tissues. However, the sequencing data do not directly establish where these siRNAs were produced or directly demonstrate RDR6-dependent amplification. The precise dosage of siRNAs mobilized from leaves that induces secondary siRNA production in cassava recipient tissues remains unclear.

Together, the results suggest that *MeAPS1a* siRNAs produced in the shoots of small cassava plants are mobile and can move from the leaves to the roots. However, root *MeAPS1a* repression was detected only in the younger SG plants. Because plant age, pot size and sampled root tissue differed between SG and LG plants, this comparison does not fully clarify the effects of plant size or transport distance. The persistence of storage root defects without detectable *MeAPS1a* repression in mature roots suggests an additional contribution from altered source metabolism, although an earlier effect of silencing on root development cannot be excluded.

### Reduction of transitory starch reduces carbon delivery to storage roots

Since *MeAPS1a* mRNA levels were not significantly reduced in the developing storage roots of LG plants **(Fig. 3C),** yet storage root agronomic parameters were strongly impacted in greenhouse-and field-grown cassava RNAi plants **(Fig. 2)**, we examined the metabolic status of storage roots more closely using the ungrafted RNAi line 61 (Ug-61) and ungrafted vector control (Ug-VC) as examples. While shoot fresh weight remained unchanged in Ug-VC and Ug-61 plants, storage root fresh weight, storage root dry matter content, and harvest index decreased significantly in Ug-61 plants **(Fig. 4A)**. Consistent with the reduction in dry matter content, the content of starch in the storage root core tissue, which consists primarily of starch-storing xylem parenchyma cells, was significantly reduced **(Fig. 4B)**. Steady state sucrose levels increased, but monosaccharide levels remained unchanged **(Fig. 4B)**. We also found that several metabolic intermediates were significantly reduced in Ug-61 plants **(Fig. S4)**. No significant differences in carbohydrates were found in the storage root peel **(Fig. 4C)**.

**Fig. 4.**
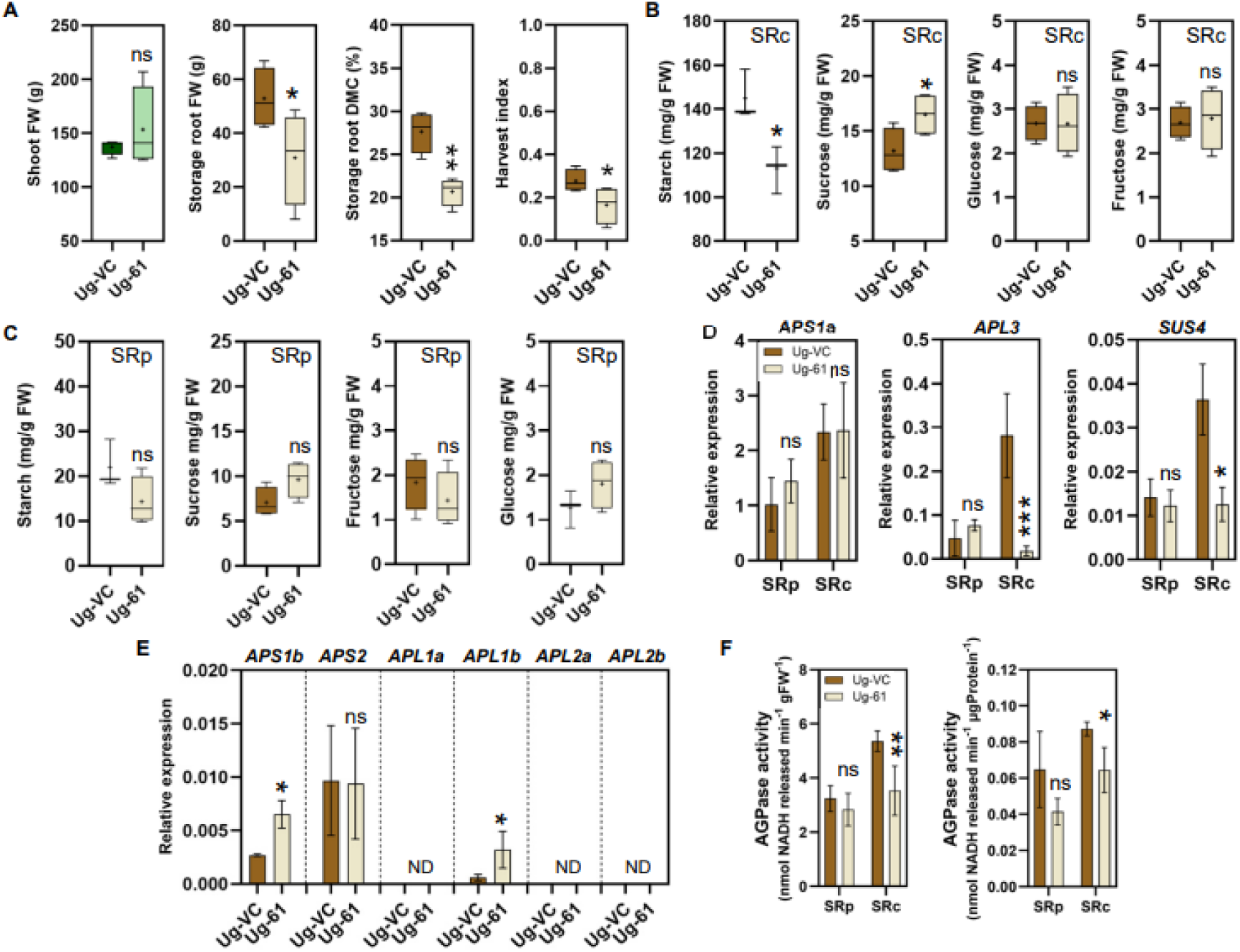
Altered storage root carbohydrate metabolism in *MeAPS1a* RNAi plants without detectable root *MeAPS1a* repression. **A)** Agronomic parameters of the harvested ungrafted VC (Ug-VC) and RNAi_61 (Ug-61) plants. **B**) Starch and soluble sugar levels in the storage root core (SRc). **C**) Starch and soluble sugar levels in the storage root peel (SRp). **D**) Transcriptional expression of *MeAPS1a, MeAPL3,* and *MeSUS4* in SRp and SRc of Ug-VC and Ug-61 plants. **E**) Transcriptional expression of other *MeAPS* and *MeAPL* genes in the SRc. **F**) AGPase enzymatic activity in SRp and SRc of Ug-VC and Ug-61 plants. Asterisks indicate the significant differences between Ug-VC and Ug-61 plants. A Student’s t-test was conducted. *, *p*<0.05; **, *p*<0.01; ***, *p*<0.005; ns, no significance. **A-F**) N = 4 plants for each Ug-VC and Ug-61.

We also analyzed the expression of the *MeAPS* and *MeAPL* genes in the peel and core of the storage roots **(Fig. 4D)**. *MeAPS1a* mRNA levels were similar in Ug-61 and Ug-VC plants. However, the expression of *MeAPL3*, the primary *MeAPL* gene variant in the storage root, was significantly lower in the core of the root. This reduction may reflect altered carbohydrate status or other developmental and metabolic changes. Sugar-responsive expression of AGPase large-subunit genes has been reported in *Arabidopsis thaliana* and potato (Müller-Röber et al., 1990; Sokolov et al., 1998). We tested *MeSUS4* as another potentially carbohydrate-regulated transcript. In potato tubers, *SUS4* is the primary tuber gene variant and is transcriptionally regulated by sucrose (Fu et al., 1995). The expression of the cassava homolog, *MeSUS4*, is dominant in storage roots and closely matches the carbohydrate status of the core root tissue (Rüscher *et al*., 2024). Similar to *MeAPL3*, the abundance of *MeSUS4* mRNA was significantly reduced in the storage root core of Ug-61 plants **(Fig. 4D)**. We examined the other *MeAPS* and *MeAPL* genes only in the storage root core tissue **(Fig. 4E)**. Unlike *MeAPL3*, *MeAPS1b* and *MeAPL1b* were upregulated in Ug-61 plants. However, their abundance was much lower than that of *MeAPS1a* and *MeAPL3*. Consequently, AGPase activity was also significantly reduced in the storage root core of Ug-61 plants **(Fig. 4F)**.

Overall, the data are consistent with reduced carbon supply from the leaves together with altered carbohydrate utilization in the storage roots. The increased steady state sucrose concentration in the root core does not directly report sucrose delivery and may rather reflect reduced sucrose utilization. These measurements therefore do not fully resolve the relative contributions of carbon delivery and sink metabolism to the reductions in storage root growth and dry matter content.

## Discussion

### Transitory starch synthesis is indispensable for cassava storage root production

Transitory starch acts as a temporary carbon reserve, sustaining plant growth and development at night and while providing physiological buffering under suboptimal or stressful conditions (Fettke and Fernie, 2024). This raises important questions for crop improvement: How much leaf starch is necessary for growth and maintenance metabolism? To what extent can “surplus” fixed carbon be redirected toward agronomically important sink organs, such as seeds, tubers, and storage roots that are important for agriculture and human consumption? Evidence from diverse species suggests that this threshold is highly species-specific, offering potential entry points for engineering carbon allocation.

In potatoes, for example, reducing AGPase activity to around 30% of wild-type levels did not hinder vegetative growth or result in nighttime carbon starvation. Instead, it promoted sucrose export during the day (Kirsten Leidreiter, 1995). Consequently, tuber yield remained unaffected (Kirsten Leidreiter, 1995; Müller-Röber et al., 1992). Furthermore, combining reduced leaf starch synthesis with enhanced sink metabolism shifted the source–sink balance in favor of tuber starch accumulation (Jonik et al., 2012).

Notably, hybrid aspen trees lacking over 95% of their transitory starch due to the knockout of two PGM genes exhibited only a 30% reduction in carbon assimilation and maintained normal growth under short-day greenhouse conditions (Wang et al., 2022). Starchless mutants of *Lotus japonicus* and *Pisum sativum* similarly displayed normal vegetative development (Harrison et al., 1998; Vriet et al., 2010). These studies suggest that some species can compensate for reduced leaf starch without experiencing significant growth impairment.

However, cassava appears to be far less tolerant of reduced transitory starch levels. Repressing *MeAPS1a* nearly eliminated leaf starch synthesis and reduced photosynthetic capacity by about 50% compared with control plants **(Fig. 1)**. Although some organ-level compensatory responses were detectable **(Fig. S2E–K)**, overall metabolism was severely impaired by the carbon shortage. Sink formation and storage root starch accumulation were markedly reduced **(Figs. 2 and 4)**. Under field conditions, RNAi plants developed smaller canopies and produced fewer and smaller storage roots **(Fig. 2R, V-X)**, resulting in substantial yield losses.

Greenhouse-grown *MeAPS1a* RNAi lines that retained small amounts of leaf starch **(line 171, Fig. 1E)** exhibited milder phenotypic defects. Notably, they had a much less pronounced reduction in photosynthesis **(Fig. 1F)**. This suggests that even low levels of leaf starch are sufficient to avoid triose-phosphate utilization limitations under greenhouse conditions. This finding also helps explain why partial repression of leaf starch in potatoes did not compromise growth. An approximately 30% residual AGPase activity likely prevented strong inhibition of photosynthesis (Jonik et al., 2012; Kirsten Leidreiter, 1995). In contrast, starch-free aspen exhibited impaired photosynthesis, yet maintained normal growth. This is presumably because its cell division rates, rather than carbon assimilation, limit overall biomass accumulation (Wang et al., 2022). Whether this conclusion holds under field conditions remains an open question.

The *MeAPS1a* RNAi cassava plants grown in the field had a more compromised canopy than the plants in the greenhouse, but they still produced storage roots with a similarly reduced yield relative to the greenhouse-grown plants **(Fig. 2J-L, V-X)**. Several mechanisms could explain why the shoot phenotype was more pronounced under field conditions. One possibility is that impaired transitory starch metabolism likely increases the risk of nocturnal carbon limitation, which affects rapidly growing shoot tissues, such as leaf primordia and expanding leaves, disproportionately. This effect may be exacerbated in the field because higher daytime assimilation rates and warmer nights increase respiratory carbon demand. Additionally, transitory starch acts as a buffer against short-term fluctuations in carbon supply, which occur more frequently in natural environments due to variable light and temperature conditions. Reducing buffering capacity could therefore destabilize the carbon supply to developing shoot tissues. In contrast, storage root bulking may depend more strongly on cumulative daytime assimilate export, making it less sensitive to short-term fluctuations in carbon availability. Overall, our results demonstrate that cassava depends heavily on transitory leaf starch for proper growth and development.

### *MeAPS1a* RNAi plants have mobile siRNAs that move systemically

Despite the availability of modern genome-editing technologies, RNA interference is still widely used because it is flexible and effective at partially repressing gene expression. Systemic movement of sRNAs has been reported in several model species, and the extent of this movement correlates with sRNA abundance in recipient tissues (Brosnan and Voinnet, 2011; Buhtz et al., 2008; Klahre et al., 2002; Voinnet and Baulcombe, 1997). However, little information exists for crop species overall.

Based on potato studies in which leaf-specific *AGPase* repression did not affect tuber development, we did not anticipate adverse effects on cassava storage roots. Therefore, to investigate carbon allocation without directly targeting storage root tissues, we expressed the *MeAPS1a* shRNA under the control of the green tissue-specific *pAtCAB1* promoter (Zierer et al., 2022) **(Fig. 1B)**.

Unexpectedly, *MeAPS1a* transcript levels were significantly lower in the storage roots of the RNAi plants, even though the shRNA itself was not detectable **(Fig. 1C)**. To test for systemic mobile siRNAs, we performed reciprocal grafting between RNAi and vector control (VC) plants and examined *MeAPS1a* expression in SG (44 DAG) and LG (127 DAG) plants **(Fig. 3)**. In the first experiment, *MeAPS1a* repression was detected in SG roots of VCx61 and 61×61 plants, but not in 61xVC plants (Fig. 3C). In the second experiment, both 154xWT and WTx154 grafts showed root repression (**Fig. S3B**). Grafting retains a basal stem segment from the rootstock, which was green when the young plants were grafted. In grafts with an RNAi rootstock, this retained tissue could provide a local source of silencing signal because *pAtCAB1* is still active in green tissues. However, siRNA production in this stem segment was not measured directly. Repression in these combinations therefore cannot be attributed solely to signals originating in the scion. The reason for the difference between the two experiments in grafts with a control rootstock remain unresolved. No *MeAPS1a* repression was detected in LG storage roots.sRNA-Seq provided direct evidence of mobile siRNAs. Twenty-to 24-nt siRNAs that perfectly matched *MeAPS1a* were present in the storage roots of both SG and LG plants, and in the leaves of LG plants **(Fig. 3)**. However, their abundance differed markedly. SG roots accumulated substantially more 20–24 nt siRNAs **(Fig. 3F)**, and their mapping profiles varied by developmental stage. SG storage roots contained siRNAs spanning the full coding sequence, whereas LG storage roots and leaves mainly contained siRNAs derived from the hairpin region **(Fig. 3G)**. This broad distribution in SG storage roots is consistent with secondary siRNA amplification, potentially involving an RDR-dependent pathway as described in Arabidopsis (Baulcombe, 2004; Taochy et al., 2017). The different mapping profiles could reflect changes in secondary siRNA production, transport or stability during development. Restricted radial movement in larger roots is another possible explanation.

Together, these findings support systemic movement of *MeAPS1a* siRNA-derived silencing signals. Lower siRNA abundance in LG roots was associated with the absence of detectable target repression. However, the experiments do not establish a silencing threshold or distinguish the effects of plant age, storage root growth, tissue composition and transport distance.

### Low carbon input severely impairs cassava storage root yield

Transitory starch repression triggered widespread metabolic disruption in cassava. Most *MeAPS1a* RNAi plants had a 50% decrease in photosynthetic capacity **(Fig. 1F)**. Sucrose levels decreased significantly, especially at night, and only slowly recovered during the day **(Fig. 1E)**. Storage root formation and starch content also decreased significantly **(Fig. 4B, C)**. Importantly, reduced storage root biomass was also observable in mature RNAi plants without any detectable root *MeAPS1a* repression (**Fig. 4**).

In mature, ungrafted RNAi plants, storage root *MeAPS1a* expression remained unchanged, while *MeSUS4* and *MeAPL3* expression and AGPase activity were reduced in the storage root core **(Fig. 4D, F)**. These changes are consistent with altered sink metabolism accompanying reduced source activity. However, sucrose-dependent regulation of *MeAPL3* was not tested directly. The increased storage root core sucrose would argue against this but could also reflect a reduced sucrose utilization, which allows sucrose to accumulate despite lower source supply. The results support a contribution of altered source-sink metabolism to the storage root phenotype, but do not establish whether downregulation of key genes in sucrose and starch metabolism is a cause or a consequence of the reduced storage root growth. The absence of detectable *MeAPS1a* repression at harvest also does not exclude a contribution from transient silencing earlier in root development.

## Conclusion

Transitory starch is essential for cassava growth and productivity. While several plant species can tolerate significant reductions in leaf starch and redirect carbon to sink organs, cassava has a narrow margin for such manipulation. Reductions in leaf starch synthesis rapidly suppress photosynthetic carbon assimilation, resulting in severe growth and storage root yield penalties. Therefore, the strong reduction of leaf starch synthesis achieved here is not a promising strategy for improving cassava source–sink relations under the conditions tested.

Our findings also underscore the importance of exercising caution when applying tissue-specific RNAi approaches in crops. Even when systemic silencing is detectable only in young plants, mobile silencing signals can complicate the interpretation of tissue-specific manipulations and their contribution to later phenotypes requires careful evaluation.

## Acknowledgments

We thank Birgit Peterson and Eva Düll for technical support with cassava tissue culture and transformation, Ingrid Schießl for cloning, and Michaela Reiser, Otilia Ciobotea, Sanika Lakshman, and David Pscheidt for technical support with experiments and greenhouse phenotyping. We also thank the NCHU CASS team field managers for the operations during the field experiment and Chih-Yun Chen, Fei-Han Chang and Ting-lun Nieh for their help during the cassava harvests.

## Author contributions

QL, WZ, SHC, WG, and US planned the experiments. QL and WZ executed laboratory and greenhouse experiments. CL generated the transgenic cassava plants. AK, JS, and QL conducted phylogenic and sRNA-seq data analysis. JH and QL performed metabolite and phytohormone measurements and analysis. TJC, SHC, and WG conducted the field experiment and analyzed the agronomic performance of the cassava lines. QL and WZ wrote and WG edited the manuscript. All authors have read the manuscript and agree with its content.

## Conflict of interest

The authors declare no conflicts of interest.

## Funding

This work was funded through grants to Prof. Uwe Sonnewald by the Bill & Melinda Gates Foundation (Grant number 008053) and Gates Agricultural Innovations (Grant number 58147). The conclusions and opinions expressed in this work are those of the author(s) alone and shall not be attributed to the Foundation. Under the grant conditions of the Foundation, a Creative Commons Attribution 4.0 License has already been assigned to the Author Accepted Manuscript version that might arise from this submission. Please note that works submitted as preprints have not undergone peer review. W.G. is supported by a Yushan Scholarship of the Ministry of Education (MOE) in Taiwan, and his research is financially supported in part by the Advanced Plant and Food Crop Biotechnology Center from The Featured Areas Research Center Program within the framework of the Higher Education Sprout Project by MOE in Taiwan.

## Data availability statement

Sequence data contained in this study is openly available in the NCBI Sequence Read Archive under BioProject accession PRJNA1481054.

## Supplementary Figures

**Fig. S1.**
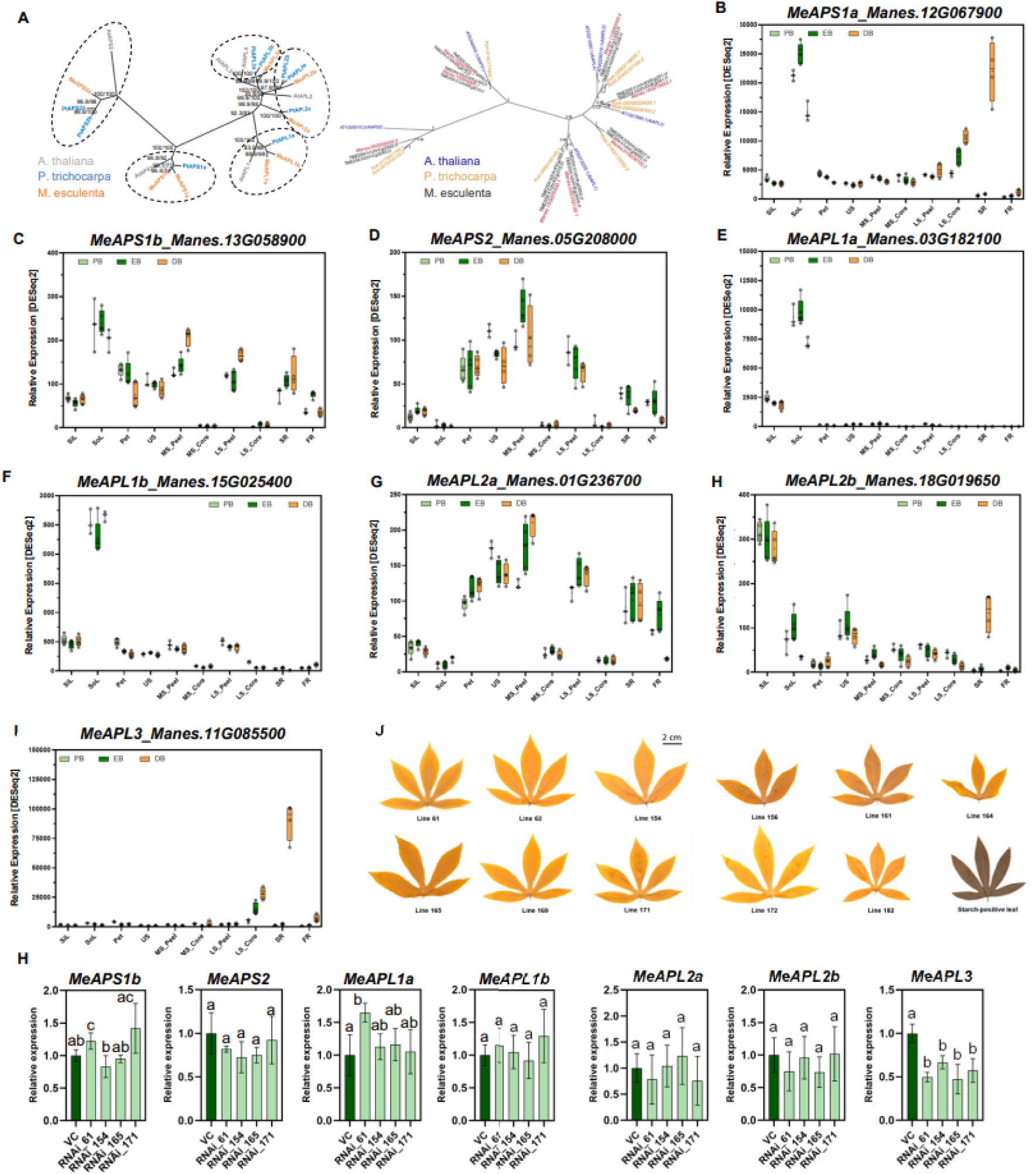
Phylogenetic and expression analysis of *MeAPS* and *MeAPL* genes. **A)** Phylogenetic analysis of *MeAPS* and *MeAPL* genes. The cassava genes in the left were obtained from the AM560 v8.1 genome assembly, and those in the right were obtained from the TME204 genome assembly (haplotype-resolved). **B**) Expression of *MeAPS1a* (*Manes*.*12G067900*). **C**) Expression of *MeAPS1b* (*Manes*.*13G058900*). **D**) Expression of *MeAPS2* (*Manes*.*05G208000*). **E**) Expression of *MeAPL1a* (*Manes*.*03G182100*). **F**) Expression of *MeAPL1b* (*Manes*.*15G025400*). **G**) Expression of *MeAPL2a* (*Manes*.*01G236700*). **H**) Expression of *MeAPL2b* (*Manes*.*18G019650*). **I**) Expression of *MeAPL3* (*Manes*.*11G085500*). PB: pre-bulking stage; EB: early-bulking stage; DB: during-bulking stage. SoL: source leaves; US: upper stem; MS_peel: middle stem peel; MS_core: middle stem core; LS_peel: lower stem peel; LS_core: lower stem core. Expression data are from Rüscher et al. (2024). **J**) Iodine staining of source leaves from 11 *MeAPS1a*_RNAi lines (yellowish and starchless), and a starch-positive leaf (dark blue) from the control plant, taken at the end of the day. **K**) Expression of *MeAPS1b, MeAPS2, MeAPL1a, MeAPL1b, MeAPL2a, MeAPL2b, MeAPL3* in source leaves of vector control (VC) plants and the selected four RNAi lines. n = 3 to 4 biological replicates. Leaf samples were collected at the midpoint of the light cycle. The letters indicate significant differences among plants based on a one-way ANOVA with Tukey’s post hoc test (*p*<0.05).

**Fig. S2.**
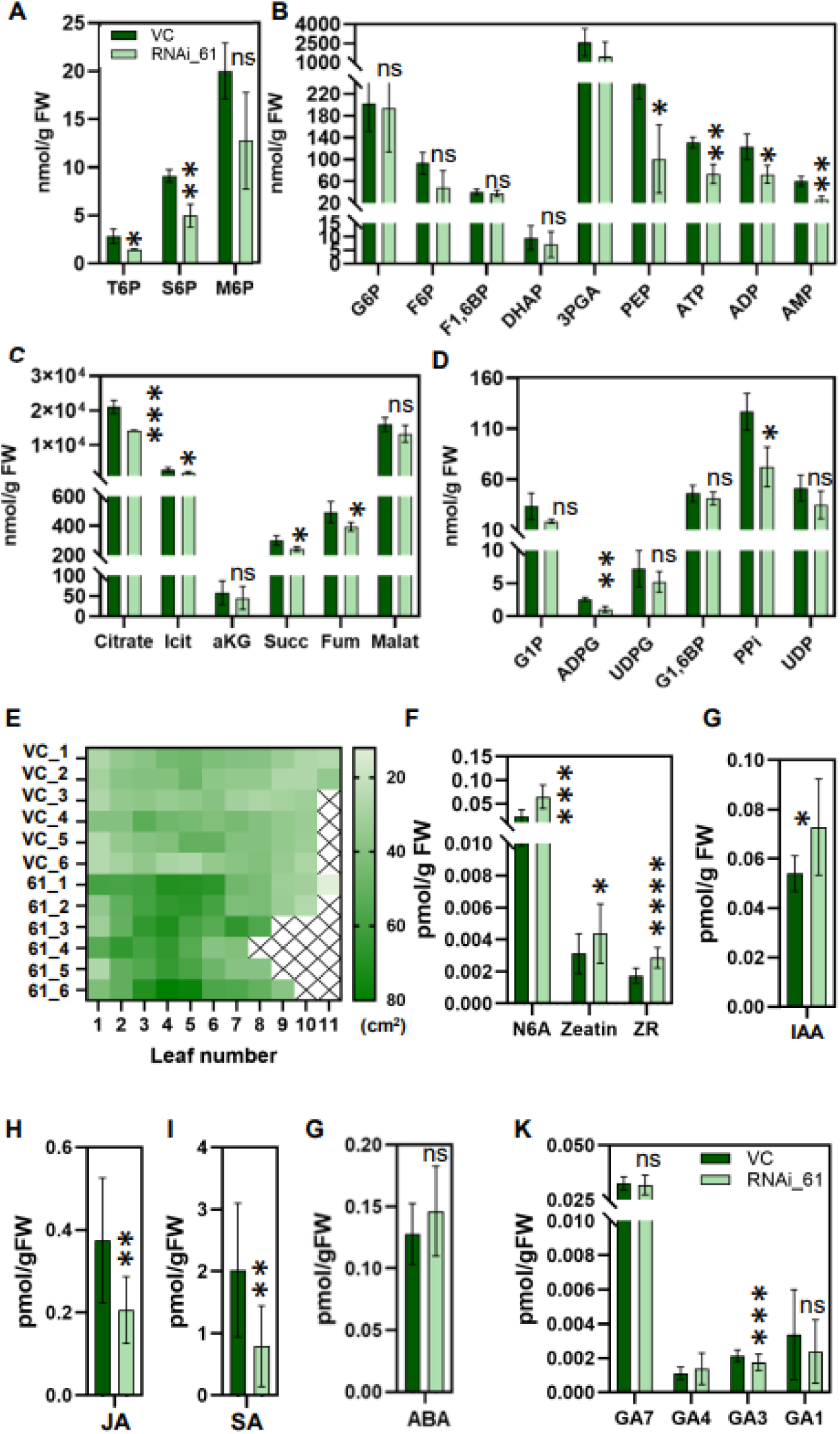
Leaf size, phytohormones, and phosphorylated metabolites levels in the leaf of the representative vector control and RNAi plants. **A-D)** Phosphorylated metabolites of the source leaves of VC and RNAi_61 plants. **E)** The size of the 1st to 11th (from apex) leaf of the VC and RNA_61 plant. N=6. **F-K)** Phytohormone levels of the 3rd-5th leaves. N=4. Asterisks indicate the significant differences; a Student’s t-test was conducted. *, *p*<0.05; **, *p*<0.01; ***, *p*<0.005; ****, *p*<0.0001; ns, no significance. The color code applies to all bar graphs.

**Fig. S3.**
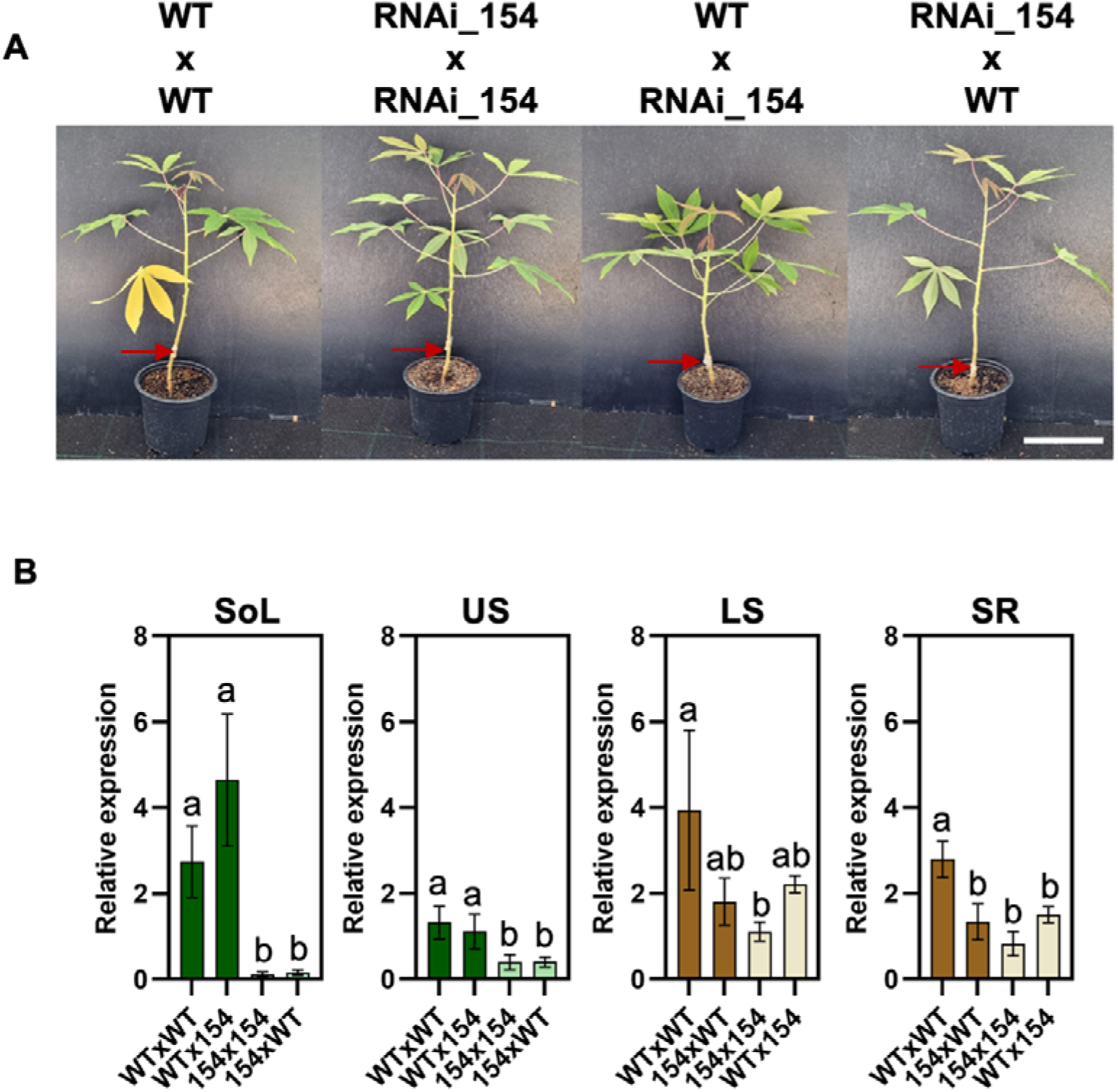
Second grafting event and evaluation. **A)** Photographs of the grafted plants of the second grafting event. Red arrowheads indicate the grafting joint. Scale bar = 10cm. **B**) Relative expression level of *MeAPS1a* in the indicated tissue of all grafts. SoL = source leaf, US = upper stem, LS = lower stem (lignified), SR = storage root. n = 3 plants. The letters in each graph indicate significant differences based on one-way ANOVA analysis with Tukey’s post hoc test (*p*<0.05).

**Fig. S4.**
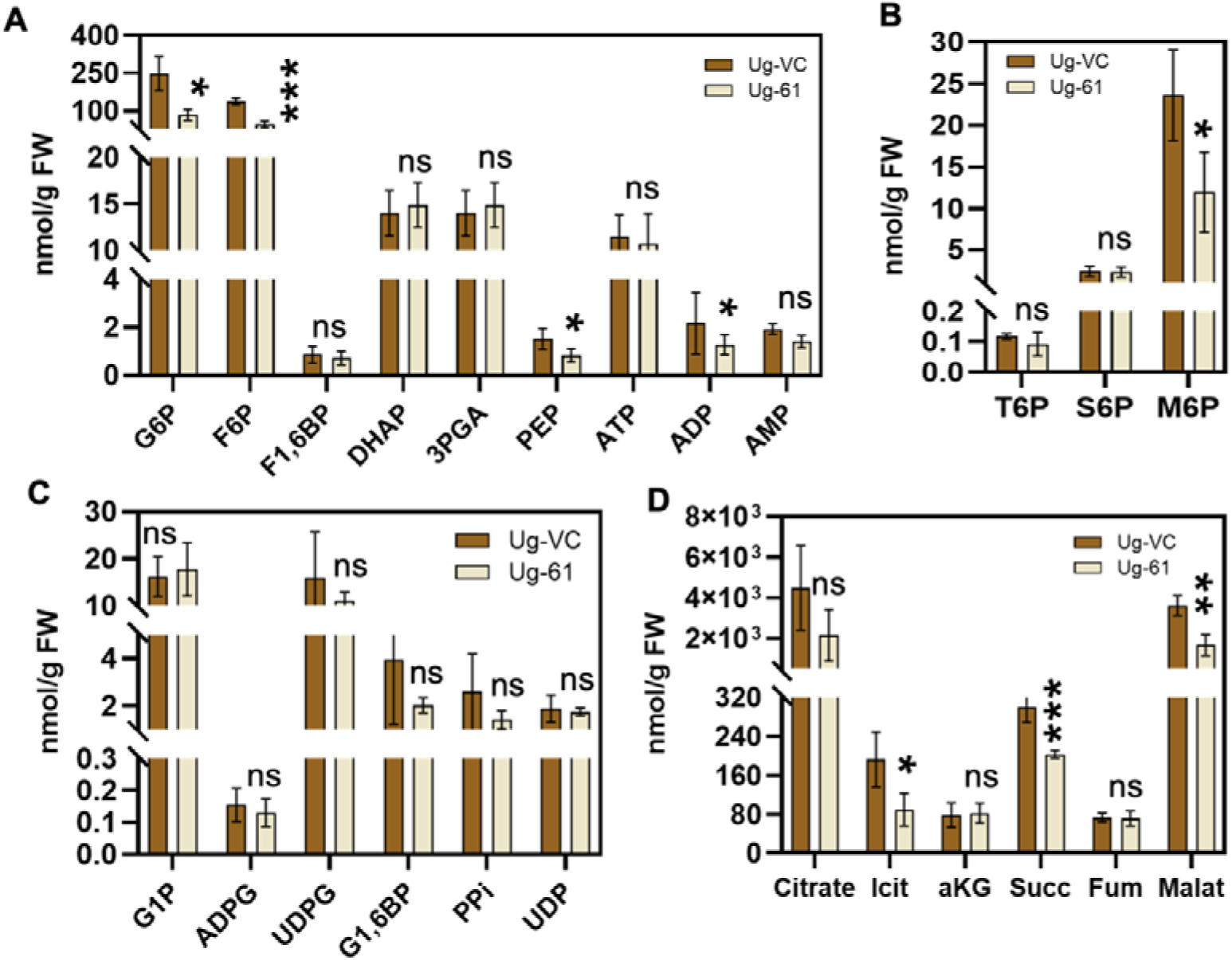
*MeAPS1a* RNAi plant still shows abnormal carbon utilization when the siRNA effect is not detectable. **A-D**) Phosphorylated metabolite profiling in SRc of Ug-VC and Ug-61. Asterisks highlight the significant difference between Ug-VC and Ug-61, a Student’s t-test was conducted. *, *p*<0.05; **, *p*<0.01; ns, no significance. **A-D**) N=4.

**Fig. S5.**
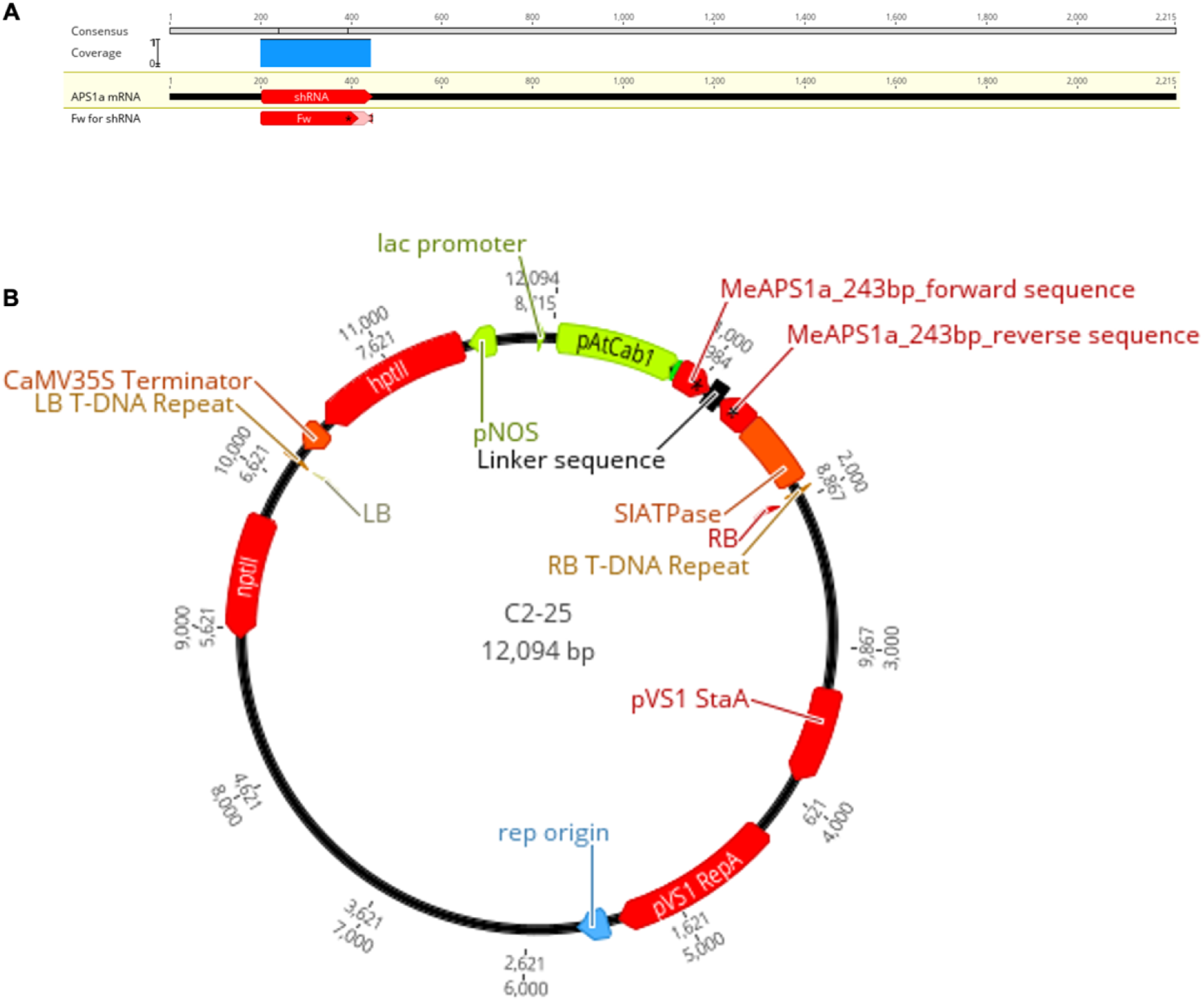
Plasmid map of the *MeAPS1a* RNAi construct. **A)** Alignment of the target sequence to the *MeAPS1a* transcript. **B**) Map of the plasmid used for *MeAPS1a*RNAi.

## Notes

### Competing Interest Statement

The authors have declared no competing interest.

